# Detection of novel allelic variations in soybean mutant population using Tilling by Sequencing

**DOI:** 10.1101/711440

**Authors:** Reneth Millas, Mary Espina, CM Sabbir Ahmed, Angelina Bernardini, Ekundayo Adeleke, Zeinab Yadegari, Korsi Dumenyo, Ali Taheri

**Author notes:** Corresponding author Correspondence to Ali Taheri.

## Abstract

One of the most important tools in genetic improvement is mutagenesis, which is a useful tool to induce genetic and phenotypic variation for trait improvement and discovery of novel genes. JTN-5203 (MG V) mutant population was generated using an induced ethyl methane sulfonate (EMS) mutagenesis and was used for detection of induced mutations in FAD2-1A and FAD2-1B genes using reverse genetics approach. Optimum concentration of EMS was used to treat 15,000 bulk JTN-5203 seeds producing 1,820 M2 population. DNA was extracted, normalized, and pooled from these individuals. Specific primers were designed from FAD2-1A and FAD2-1B genes that are involved in the fatty acid biosynthesis pathway for further analysis using next-generation sequencing. High throughput mutation discovery through TILLING-by-Sequencing approach was used to detect novel allelic variations in this population. Several mutations and allelic variations with high impacts were detected for FAD2-1A and FAD2-1B. This includes GC to AT transition mutations in FAD2-1A (20%) and FAD2-1B (69%). Mutation density for this population is estimated to be about 1/136kb. Through mutagenesis and high-throughput sequencing technologies, novel alleles underlying the mutations observed in mutants with reduced polyunsaturated fatty acids will be identified, and these mutants can be further used in breeding soybean lines with improved fatty acid profile, thereby developing heart-healthy-soybeans.

## INTRODUCTION

Soybean (*Glycine max* L. Merrill) is an oil-producing crop under the legume family, Fabaceae, and constitute over 33% of the total area planted for crops in the US alone. Soybean oil comprised 61% of the world oilseed production, and 55% of the US vegetable oil consumption (SoyStats, 2018). The fatty acid profile of the soybean determines its utilization and applications. From the five major fatty acids in the soybean oil, α-linolenic acid, being an unstable component, is responsible for the poor characteristics of soybean such as undesirable odor and reduced storage life ^2,3^. To reduce the level of linolenic acid, soybean is usually subjected to partial hydrogenation that results in the formation of trans fatty acids that are linked to coronary heart disease ^4^. In order to improve the stability and shelf life of soybean oil without hydrogenation, soybean with increased oleic acid content should be developed and characterized through mutation breeding.

Molecular breeding approaches used to alter the seed oil composition are targeting important genes involved in the pathway leading to fatty acid biosynthesis in seed. In the biosynthesis pathway, the oleic acid (C18:1) undergoes desaturation to linoleic acid (C18:2) by the action of microsomal enzymes delta-12-fatty acid desaturase (FAD2)^5^. Five FAD2 members, which were observed in four different loci, constitute the fatty acid desaturase family in the soybean genome ^6–9^. Two microsomal FAD2-1 desaturases, i.e. *FAD2-1A* (*Glyma10g42470*) and *FAD2-1B* (*Glyma20g24530)* are primarily expressed in developing seeds. The FAD2 genes in soybean determine the levels of monounsaturated fats in soybean oil and vegetative tissues^10,6.^

One of the most important non-genetically-modified (GM) tools in crop improvement is mutagenesis, which results in the introduction of genetic variation and occasionally generating mutants with improved traits or novel phenotypes ^11,12^. For breeding purposes, chemical mutagenesis and irradiation are the established methods that have been utilized to generate mutant plants ^13,14^. Chemical mutagenesis, either with EMS (Ethyl methanesulfonate) or NMU (Nnitroso-N-methylurea), usually causes single nucleotide polymorphisms that are greatly important for studying gene function as well as for their potential use in crop improvement. Among the chemical mutagens, EMS is used frequently because it creates a high frequency of non-lethal point mutations ^15^. EMS mutagenesis can induce changes in the gene of interest and can be used to study the gene functions using reverse genetics approach ^16^.

To detect the nucleotide changes in the genome of the mutant populations, molecular screening methods have been developed including Targeting Induced Local Lesions IN Genomes (TILLING) and TILLING-by-Sequencing (TbyS) ^14,17^. TILLING is a high-throughput reverse genetics technique used for identifying novel mutant alleles from mutagenized populations ^18–20^ and to obtain allelic series from a chemically mutagenized population ^16^. It is also a method used to identify unique, chemically-induced mutations within the target genes which can alter the gene expression and functions ^19,21^. By using this technique, more alleles can easily be generated at a target locus, and elite mutant alleles can be readily available in conventional breeding programs^22,23^. It also allows for identification of novel variation in target genes which is important for developing breeding germplasm with improved traits of interest. TILLING depends mainly on the capacity of a mutagen, which is most commonly the chemical mutagen ethyl methanesulfonate, to generate point mutations that produce novel SNP across the genome of each mutant line ^24^. As a non-transgenic reverse genetic approach, TILLING has been successfully used to identify mutations in genes controlling important agronomic traits. In the legume species, *Lotus japonicus* was first used to screen for mutations using the TILLING platform ^12^. So far, two soybean cultivars including Williams 82 and Forrest were chemically mutagenized and the TILLING protocol was successfully applied ^16^. However, this technique has limitations in studying soybean traits such as oil composition due to the gene copy number and similarities in the soybean genome ^25^.

With the reduced cost in sequencing technologies, TbyS has been developed and utilized for the identification and detection of such induced or natural mutations like point mutation, insertion and deletion ^14,17^, and to discover rare mutations in the population. This can also be used to identify and characterize the genes controlling the trait of interest within the mutant populations ^26^. TbyS is the application of high-throughput sequencing technologies coupled with multidimensional pooling, and bioinformatics pipeline that results to an efficient detection of allelic variation in mutant populations. TbyS provides high sensitivity and specificity, and is more effective than TILLING. TbyS discovery was first reported in rice and wheat using either bi- or tridimensional pooling schemes ^27^. Thus, TbyS allows the identification of plant lines in which a mutation has been successfully induced in the target gene of interest.

Due to the increasing demand of high oleic soybeans, recent study has aimed to produce mutant lines with high oleic acid content. This study will utilize the EMS-induced mutant population for detection of induced mutations in the FAD2-1A and FAD2-1B genes using TILLING-by-Sequencing approach.

## MATERIALS AND METHODS

### Mutant generation, DNA extraction, and pooling

The development and generation of JTN-5203 mutant population as well as the DNA extraction and normalization are described earlier ^28^. The DNA of 6,400 individual M2 mutants were used to perform 2-rounds of pooling to generate a TILLING population. From the 67 boxes comprising the 6,400 individual DNA samples, 75 ul each of 50ng DNAs were pooled into 8 boxes, which constitute the first-round of pooled DNA. The first pooled DNAs were then pooled to generate one box that constitute the second-round final pooled DNA and was used as working DNA template for high-throughput sequencing.

### Gene-specific primer design

Gene-specific primers amplifying the Fatty Acid Desaturase 2 (*FAD2-1A* and *FAD2-1B*) genes were designed using PCR Tiler v1.42 with a built-in specificity check for *Glycine max* ^29^. Tables 1 and 2 show the details of eight FAD2-1A and 10 FAD2-1B primer pairs designed for Illumina sequencing. These primers were designed to cover the whole gene plus 400 upstream and downstream of each gene. Illumina adapters were attached to the 5’ end of each primer in order to prepare the amplicon primers for Illumina sequencing. The designed primers were ordered and synthesized by Thermo Fisher Scientific.

**Table 1.**
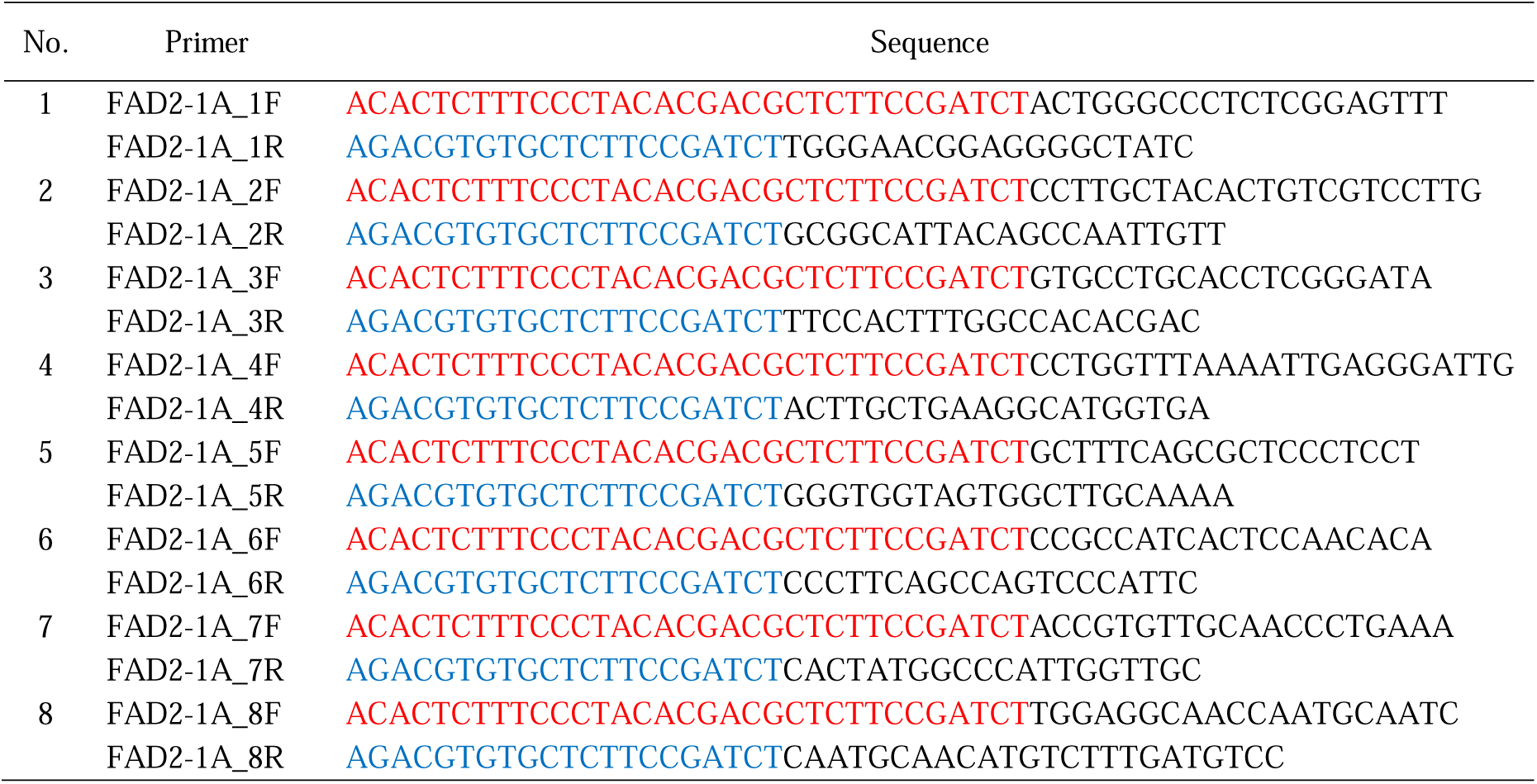
FAD2-1A gene-specific primers for Illumina sequencing.

**Table 2.**
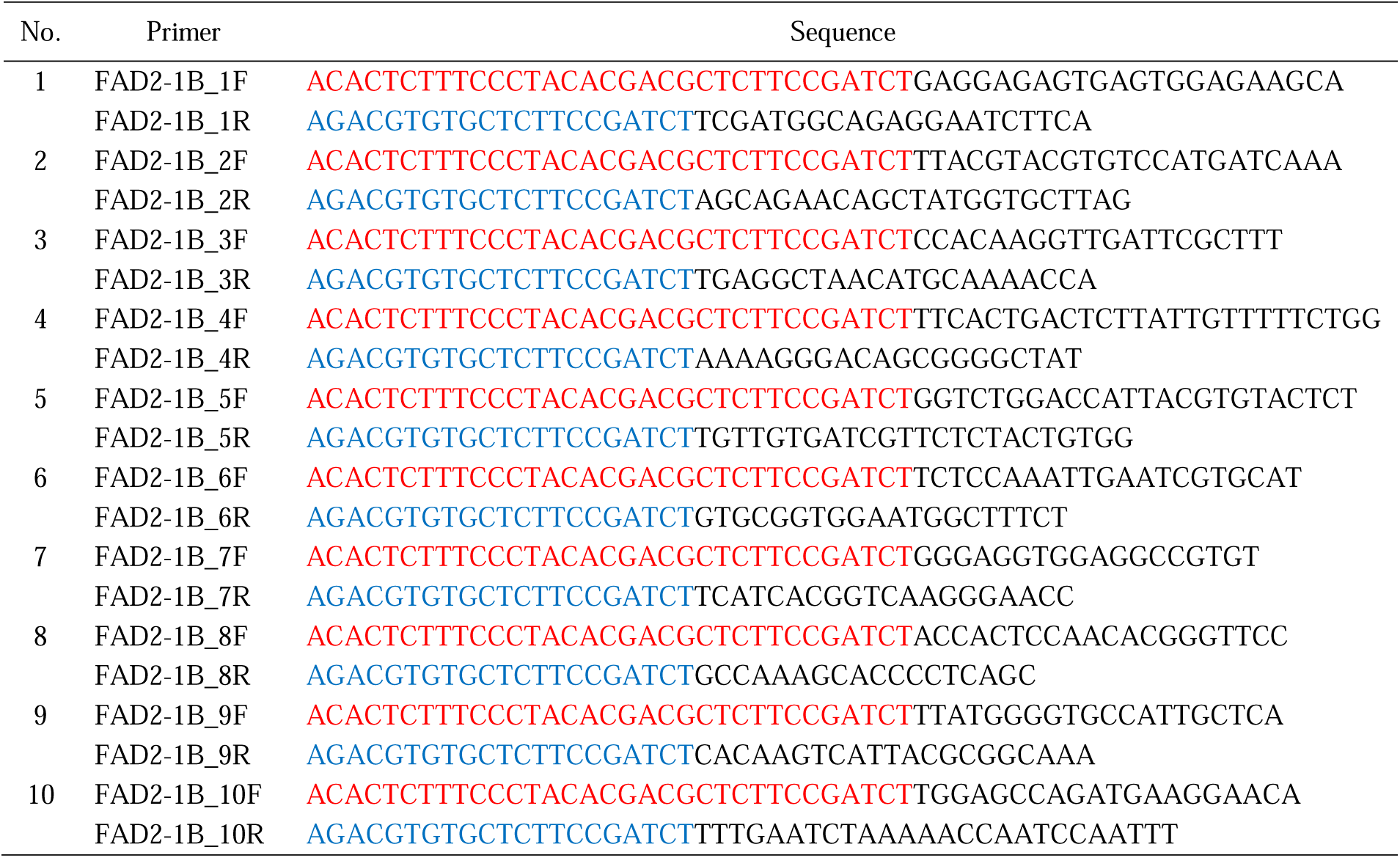
FAD2-1B gene-specific primers for Illumina sequencing.

### Dual-index library preparation

The workflow of MiSeq dual indexing library preparation involved the following steps: *Inner PCR, Clean-up#1, Outer PCR, Pre-pooling, Clean-up#2*, and *Final pooling*. Inner PCR with the tailed primers was performed using a high-fidelity enzyme mix to amplify the region of interest from the genomic DNA. These comprised of eight individual PCRs for FAD2-1A and 10 individual PCRs for FAD2-1B. The total PCR reaction volume was 30 μl, containing 8.57 μl 2X KAPA HiFi HotStart ReadyMix (Kapa Biosystems, Wilmington, Massachusetts), 1.29 μl of each primer (10 μM), 14.57 μl sterile DNA-free water, and 4.28 μl 50ng DNA. The amplifications were performed in a Bio-Rad Gradient Thermal Cycler (Hercules, CA) under the following conditions: initial denaturation of 2 mins at 98°C, followed by 27 cycles of run with denaturation at 98°C for 20 s, annealing at 64°C for 30 s, and extension at 72°C for 1 min. A single cycle of 2 mins at 72°C for extension was provided at the end of amplification reactions. The amplified products were verified by electrophoresis (Bio-Rad, Hercules, CA) using 1.5% agarose at 80V for 40 minutes. Then, a *Clean-up#1* step was performed for all the individual PCR products to remove loose primers, primer dimers, and unspecific products using AMPure magnetic beads and following the manufacturer’s protocol.

Next, the outer PCR step was carried out to barcode the cleaned amplicon product with dual indexes. The P7 and P5 indexed primers contained the Illumina sequencing handles, which allowed the barcoded DNA fragments to attach to the Illumina flow cell surface during sequencing (Table 3). The total PCR reaction volume was 26 μL, containing 12 μl of cleaned PCR product, 14 μl 2X KAPA HiFi HotStart ReadyMix, and 1 μl of each primer (10 μM). The amplifications were performed in a C1000 Bio-Rad Gradient Thermal Cycler (Hercules, CA) under the following conditions: initial denaturation of 2 mins at 98°C, followed by 9 cycles of run with denaturation at 98°C for 20 s, annealing at 62°C for 30 s, and extension at 72°C for 30 s. A single cycle of 2 mins at 72°C for extension was provided at the end of the amplification reactions. The amplified products were verified by gel electrophoresis (Bio-Rad, Hercules, CA) using 1.5% agarose at 80V for 35 minutes. Next, pre-pooling was done for the outer PCR products to generate one final PCR plate. This was performed by measuring the DNA concentration of samples, and barcoded amplicons were pooled at equal mass (for instance, pool barcoded amplicons from the same column in the plate at 5ul each). Then, a second clean-up was performed by using AMPure magnetic beads and following the manufacturer’s protocol. Lastly, the concentration of cleaned pre-pooled samples was measured and samples were pooled in equal molar amount to generate the final library for sequencing. The final library was verified by gel electrophoresis (Bio-Rad, Hercules, CA) using 2% agarose with 1 kb ladder at 80V for 45 minutes.

**Table 3.**
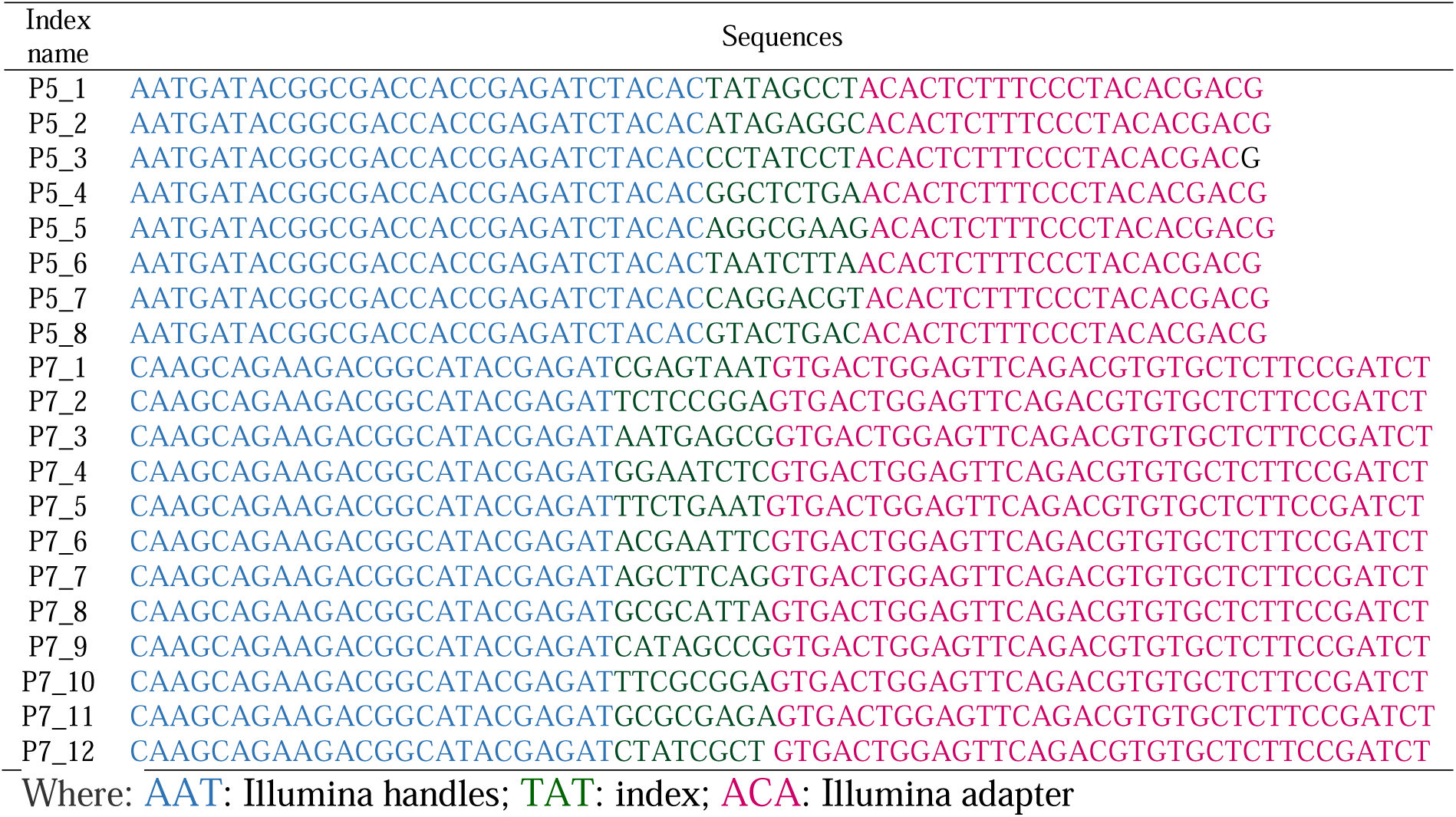
Detailed sequences of the index primers for Illumina sequencing.

### Illumina sequencing and analysis

Final library (18 ng/ul, 50ul) was submitted to the sequencing center at Vanderbilt University for high-throughput sequencing using Illumina Miseq (v2, PE 300 cycle) with the paired-end multiplexed library. Trimmomatic (Version 0.36) ^30^ was used for trimming the sequencing adapters and low quality reads from the raw data, and HISAT2 ^31^ was used to align the reads to the reference genes (*Glyma*.*10G278000* for FAD2-1A and *Glyma20g24530* for FAD2-1B). Samtools mpileup ^32^ and VarScan2 ^33^ were used to call the variants at a single base pair resolution.

## RESULTS AND DISCUSSION

A new EMS-induced soybean mutant population, JTN-5203, was developed by using EMS at 60 Mm concentration and used for high throughput screening of targeted lesions at FAD2-1A and FAD2-1B genes. A total of 6,400 individual M2 mutants were generated and DNA was extracted from each plant. Pooling of DNA was then performed to generate a TILLING population. From a total of 67 boxes comprising 6,400 individual DNA, two rounds of pooling were carried out to generate one box that constitute the final pooled DNA and was used as template for detection of the induced mutations through TbyS approach.

To ensure good coverage of the sequencing results in the genome, 400 bp additional sequences were added on both upstream and downstream regions of the FAD2-1A and FAD2-1B genes. Hence, in this experiment, the size of the FAD2-1A gene used was 2,777 bp and 4,234 bp for FAD2-1B. To detect the mutations that were present in the population, amplicon sequencing using dual index barcoding was employed in the study. Eight individual PCRs for FAD2-1A and 10 PCRs for FAD2-1B were prepared and used to perform the dual indexed library preparation steps that include inner PCR, clean-up 1, outer PCR, pre-pooling, clean-up 2, and final pooling. Final FAD2-1A and FAD2-1B libraries were generated and submitted to Vanderbilt University for Illumina sequencing. IGV was used to view the gene structure and the type and distribution of the SNPs that were detected using TbyS for FAD2-1A (Figure 1) and FAD2-1B (Figure 2).

**Figure 1.**
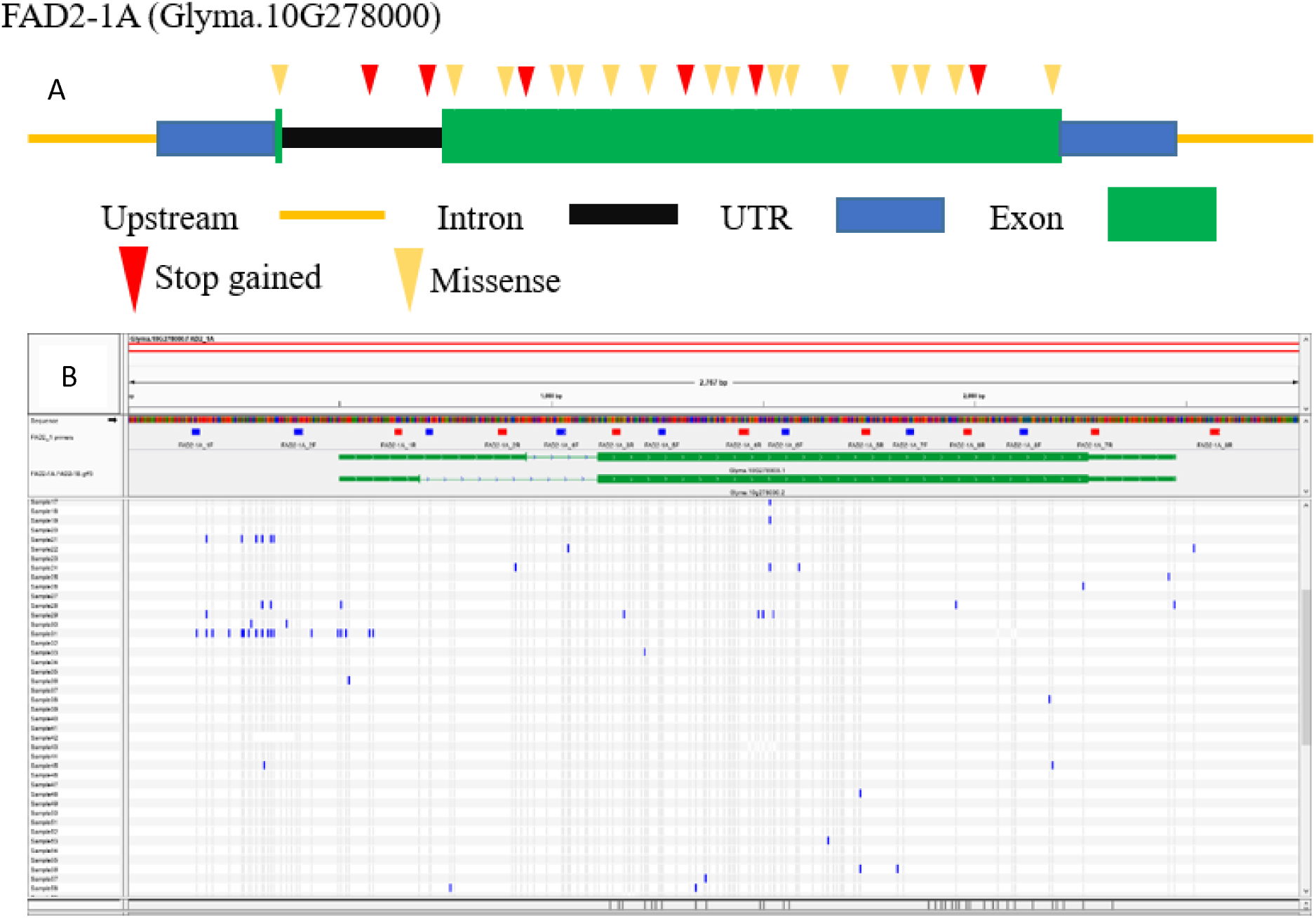
A) Type and distribution of induced mutations discovered in FAD2-1A detected using TbyS. Synonymous and mutations in UTR regions are not displayed in this diagram. B) Location of primer pairs spanning on the gene (Blue = Forward and Red= Reverse primers).

**Figure 2.**
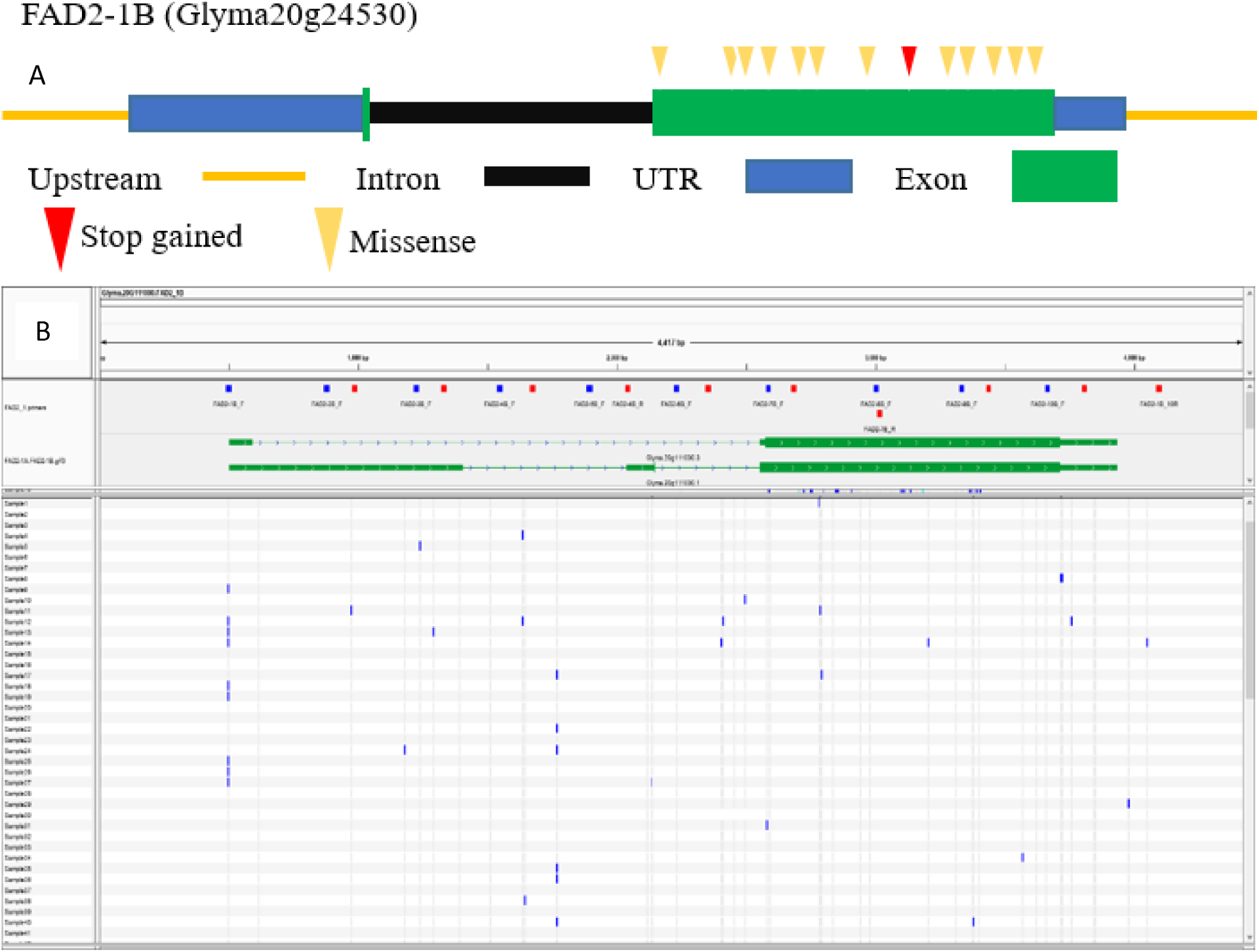
A) FAD2-1B gene structure and some of the SNPs that were detected using TbyS. Synonymous and mutations in UTR regions are not displayed in this diagram. B) Location of primer pairs spanning on the gene (Blue = Forward and Red= Reverse primers).

Figure 3 displays the variations and the effects of mutations that were detected in the TbyS for both the entire gene and coding region of FAD2-1A. The result shows that the consequence of 73% of mutations that were detected in the whole genome led to an upstream gene variant, which means that nucleotide polymorphisms are located mostly in the 5’ region of the gene. In addition, several other mutations in FAD2-1A gene lead to missense variant (14%), intron variant (4%), synonymous variant (4%), stop gained (2%), downstream gene variant (1%), 5’ UTR variant (1%), and splice region variant (1%). A total of 151 induced mutations were detected in the FAD2-1A coding region, most of these mutations resulted in a missense variant (70%), synonymous variant (21%), stop gained (8%), and start lost (1%). Regarding the impact, 8.61% of the mutations had high impact (e.g., stop gained and start lost), 21.19 % had low impact (e.g., synonymous variant), and majority of the mutations (70.20%) had a moderate impact (e.g., missense mutation) on the gene function. Of the 13 high-impact mutations, two mutations resulted to a stop gained. These mutants carried GC to AT transition that are in agreement with the expected base changes for an EMS-induced mutation. To further look at the induced point mutations that were detected in the mutants, the spectrum of mutations that were sequenced at the FAD2-1A gene is presented in Table 4. The pipeline predicted 151 mutations after screening 6400 individual M2 plants for FAD2-1A (2428bp including intron and 400bp up- and down-stream region). The mutation density was estimated as the total number of mutations divided by the total number of base pairs screened (amplicon size × individuals screened) (5) and was about ∼1/136kb. Earlier reports has indicated a mutation frequency of 1/140-550 using 40mM EMS concentration in soybean populations ^16^. It has been reported that EMS mutagenesis induces G/C to A/T transitions most of the time ^16^. However, only 20% of the observed mutations in this study were G/C to A/T transitions. Hence, most conservative estimation of mutation frequency will only consider such transitions and the mutation density will be ∼1/700kb in this population.

**Table 4.** The spectrum of induced point mutations that were sequenced at the FAD2-1A and FAD2-1B genes.

| Gene | G→A | C→T | A→G | T→C | A→T | T→A | C→A | C→G | G→C | G→T | T→G | A→C | Total |
| --- | --- | --- | --- | --- | --- | --- | --- | --- | --- | --- | --- | --- | --- |
| FAD2-1A | 16 | 8 | 4 | 4 | 4 | 6 | 38 | 4 | 2 | 44 | 21 | 0 | 151 |
| FAD2-1B | 35 | 42 | 0 | 0 | 7 | 7 | 7 | 0 | 0 | 0 | 7 | 7 | 112 |

**Figure 3.**
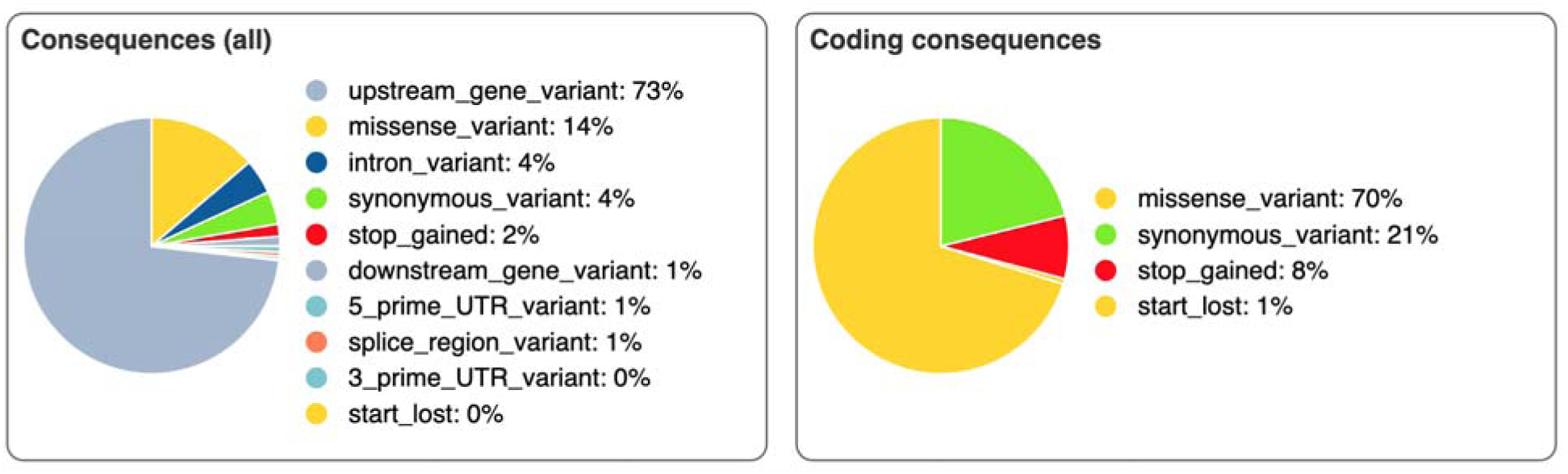
Consequences of the mutations that were detected for both the whole gene and coding region for FAD2-1A gene.

For FAD2-1B gene, the effects of the mutations for both the whole gene and coding regions is shown in Figure 4. The mutations in the complete gene include, upstream gene variant (36%), missense variant (21%), 5’ UTR variant (14%), 3’ UTR variant (9%), intron variant (9%), synonymous variant (5%), downstream gene variant (5%), and stop gained (2%). The results could mean that several random mutations were widely spread across the genome. A total of 112 mutations were detected in the FAD2-1B coding region, and most of these mutations resulted in a missense variant (75%), synonymous variant (19%), and stop gained (6%). In terms of their impact, 6.25% of the mutation had a high impact (e.g., stop gained), 18.75% had low impact (e.g., synonymous variant), and majority (75%) had moderate impact (e.g., missense mutation) on the gene function. Interestingly, all of the mutations with high and low impact on the FAD2-1B gene function carried a GC to AT transition that are in agreement to the expected mutation for an EMS-induced mutant. Further, the spectrum of induced point mutations for FAD2-1B gene is also presented in Table 4. The result shows that 69% of the identified mutations conformed to the predicted GC to AT transitions of the EMS-induced mutations. However, transversion mutations (TA to AT, CA to AC, and GT to GT) were also detected in the EMS-induced population.

**Figure 4.**
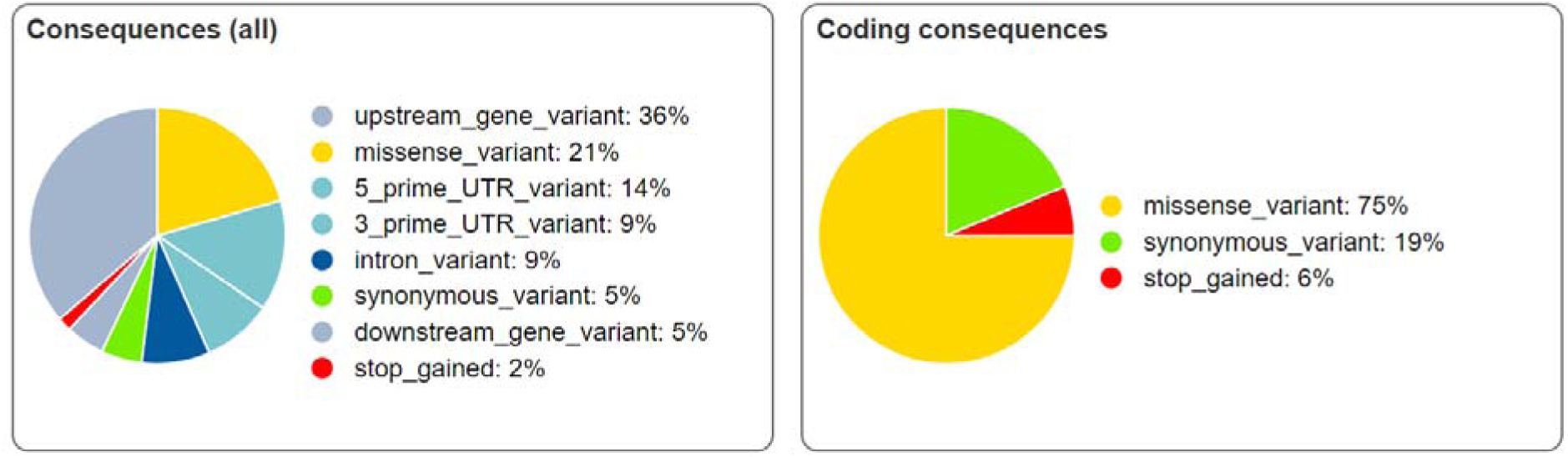
Consequences of the mutations that were detected for both the whole gene and coding region for FAD2-1B gene.

Majority of the mutations, especially with FAD2-1B gene, are in accordance with the predicted G/C to A/T transitions for an EMS induced mutants, and in agreement with the nucleotide substitutions observed in the previous study of EMS-mutagenized soybean ^16^. The results imply that the use of high throughput mutation discovery through TILLING-by-Sequencing approach has been successfully applied to the new EMS-induced soybean mutant population. However, transversions were also detected for both FAD2-1A and FAD2-1B genes, and could be false positives. However, similar findings were also observed in other studies for soybean ^16^, rice ^11^, tomato ^34^, barley ^35^, and squash ^36^. These transversion mutations maybe caused by unknown mechanisms, but are still produced by EMS ^36^. In addition, these could also be random point mutations and other low level chromosomal breaks and lesions ^37^. The transversion mutations that were detected could also be mutation biases and effect of genotype background.

The mutations that were detected are useful as basis for identification of specific pools that contain mutants that have a novel genotypic variation and could ultimately be used for screening for high oleic phenotype and also for other plant breeding purposes.

## CONCLUSION

An EMS-induced soybean mutant population was developed and used for detection of induced mutations at FAD2-1A and FAD2-1B genes using reverse genetics approach. The high throughput mutation discovery through TILLING-by-Sequencing approach has been successfully applied to the new EMS-induced soybean mutant population. Novel allelic variations in FAD2-1 and FAD2-1B genes were observed. Both FAD2-1A and FAD2-1B carried mutations that have several impacts on gene function. These include GC to AT transition that were consist of 20% in FAD2-1A and 69% in FAD2-1B. These mutations confirmed to the predicted GC to AT transitions of the EMS-induced mutations. The mutations that were detected are useful for identification of specific pools that contain mutants that have a novel genotypic variation and could ultimately be used for screening for high oleic phenotype and also other plant breeding purposes.

## Author contributions

R.M. drafted the manuscript, performed DNA pooling, library prep and Tilling by Sequencing. M.J.E., S.A., A.B., E.A., Z.Y. developed the EMS population and extracted DNA from this population. K.D and A.T. edited the manuscript and conceived the project, designed, and planned the experiment. K.D and A.T supervised students and oversea the work.

## Competing Interests

The authors declare no competing interests.

